# RUNX1-ETO expression in epidermal keratinocytes induces progressive skin inflammation *in vivo*

**DOI:** 10.64898/2026.05.10.724156

**Authors:** Hinata Date, Mizuho Ishikawa, Ikuto Nishikawa, Hung Manh Phung, Nguyen Thi Kim Nguyen, Goro Sashida, Motomi Osato, Aiko Sada

**Affiliations:** Laboratory of Skin Regeneration and Aging, International Research Center for Medical Sciences (IRCMS), Kumamoto University, Kumamoto, Japan; Division of Skin Regeneration and Aging, Medical Institute of Bioregulation, Kyushu University, Fukuoka, Japan; Laboratory of Transcriptional Regulation in Leukemogenesis, International Research Center for Medical Sciences (IRCMS), Kumamoto University, Kumamoto, Japan; Department of Medical Technology, Kumamoto Health Science University, Kumamoto, Japan

**Keywords:** Skin inflammation, Epidermal keratinocytes, RUNX1-ETO, Inflammatory pathways

## Abstract

Basal keratinocytes in the skin are essential for epidermal homeostasis and repair; however, how intrinsic alterations in these cells contribute to inflammatory skin pathology remains poorly understood. In this study, we employed a tamoxifen-inducible mouse model to express the human RUNX1-ETO fusion gene, a well-established oncogenic driver of acute myeloid leukemia, in epidermal basal keratinocytes. RUNX1-ETO induction in keratinocytes resulted in progressive skin inflammation *in vivo*, accompanied by splenomegaly, epidermal hyperplasia, increased cytokine production, and alterations in epidermal stem cell composition. Inflammatory lesions were prominent in the tail, ear, and plantar epidermis, whereas hair-bearing dorsal skin remained largely unaffected. RNA-seq analysis of FACS-isolated RUNX1-ETO^+^ basal keratinocytes revealed global changes in gene expression, characterized by the suppression of epidermal homeostatic and metabolic programs and the activation of inflammatory signaling pathways. In particular, RUNX1-ETO expression was associated with increased TNF/NF-κB and IL-6–STAT signaling, as well as interferon-associated inflammatory pathways, together with the induction of neutrophil-attracting chemokines and epithelial inflammatory mediators. Together, these findings indicate that RUNX1-ETO-mediated transcriptional dysregulation in basal keratinocytes promotes a pro-inflammatory cellular state that drives progressive skin inflammation.

## Introduction

The interfollicular epidermis serves as the outermost barrier and plays a central role in defenses against environmental stimuli and infection. Beyond their role in maintaining tissue structural integrity, epidermal keratinocytes actively contribute to regulating skin immunity (Simmons and Gallo, 2024). Because the epidermis is constantly exposed to various stimuli, keratinocytes function as sentinels that detect microbial and danger signals and produce antimicrobial peptides, cytokines, and chemokines that coordinate immune responses and tissue repair (Ye and Lai, 2025). Dysregulation of these epidermal programs has been implicated in inflammatory skin diseases. For example, activation of the transcription factor Stat3 in epidermal keratinocytes drives psoriasis-like lesions (Ohtani et al., 2000). Transcription factors such as Stat3 and AP-1 can establish persistent chromatin states in epidermal stem cells, conferring inflammatory memory that enhances subsequent inflammatory responses (Larsen et al., 2021; Naik et al., 2017). These findings suggest an active role for epidermal keratinocytes in coordinating skin immune responses; however, the keratinocyte-intrinsic mechanisms that perturb homeostatic transcriptional programs and drive pathological inflammatory activation in the epidermis remain poorly understood.

Runx1 (also known as AML1), a transcription factor discovered in 1991 in an acute myeloid leukemia (AML) patient (Miyoshi et al., 1991), plays essential roles in hematopoiesis and leukemia development (Lacaud et al., 2002). Chromosomal translocations involving RUNX1, such as t(8;21)(q22;q22), generate the RUNX1-RUNX1T1 fusion gene (also known as RUNX1-ETO), an oncogenic driver of AML (Reikvam et al., 2011). RUNX1-ETO primarily acts as a dominant-negative inhibitor, interfering with RUNX1-associated functions and perturbing gene regulatory states (Gelmetti et al., 1998; Mulloy et al., 2002; Gardini et al., 2008). In addition to inhibiting RUNX1-dependent transcription, RUNX1-ETO induces epigenetic remodeling by altering chromatin accessibility and transcription factor binding (Ptasinska et al., 2012). Thus, although RUNX1-ETO is primarily studied in hematopoietic malignancy, it provides a useful experimental system to examine how the perturbation of RUNX1-associated cellular regulatory programs affects tissue homeostasis *in vivo*.

In the skin, Runx1 has been shown to regulate hair follicle stem cell function and fate and contribute to epithelial tumor formation by enhancing the JAK–STAT pathway (Hoi et al., 2010; Scheitz et al., 2012). Runx1 also orchestrates stem cell–niche interactions and regulates hair cycle progression by modulating the cell cycle regulators and lipid metabolism that control the proliferative state of hair follicle stem cells (Lee et al., 2013; Jain et al., 2018; Li et al., 2019). Runx1 expression is tightly regulated during the hair cycle, dynamically shaping differentiation and apoptotic programs in hair follicle stem cells (Lee et al., 2018). These findings illustrate the multifunctional, context-dependent roles of Runx1 in regulating hair follicle stem cells and in tumorigenesis (Scheitz and Tumbar, 2013). Despite these insights, the role of Runx1 in the interfollicular epidermis remains poorly understood.

RUNX1 has also been implicated in human skin pathology. AML patients carrying RUNX1 mutations develop leukemic skin lesions (Rao and Danturty, 2012). Germline RUNX1 mutations causing familial platelet disorder (FPD) are associated not only with thrombocytopenia and a predisposition to myeloid malignancy but also with abnormal inflammatory responses and psoriasis-like skin manifestations (Brown et al., 2020). Psoriasis-associated single-nucleotide polymorphisms that disrupt RUNX1 binding sites have also been identified as genetic susceptibility factors for psoriasis (Helms et al., 2003; Harden et al., 2015). Runx1 also regulates T cell differentiation and effector functions (Taniuchi et al., 2002; Mallone et al., 2011), and RUNX1 haploinsufficiency can increase STAT3 activation through hypersensitivity to G-CSF (Chin et al., 2016). Together, these findings suggest that RUNX1 contributes to immune regulatory pathways that may influence skin inflammatory conditions.

Previous studies have described skin abnormalities in RUNX1-ETO/eR1-CreER transgenic mice, driven by the Runx1 enhancer (Abdallah et al., 2021). However, the involvement of skin epidermal keratinocytes in this abnormality has not been clarified. In the current study, we established a mouse model with epidermal-keratinocyte-specific RUNX1-ETO induction to determine the effects of RUNX1-ETO on epidermal homeostasis and skin inflammation. RUNX1-ETO induction disrupted normal epidermal keratinocyte function, promoting a shift toward the pro-inflammatory state and leading to progressive skin inflammation. These results establish RUNX1-ETO as a keratinocyte-intrinsic model to study epidermal–immune crosstalk, with RUNX1-ETO playing a unique role in the initiation or progression of inflammation-prone keratinocyte states observed in inflammatory skin diseases.

## Results

### Induction of RUNX1-ETO in epidermal keratinocytes leads to progressive skin inflammation

In a previous study using RUNX1-ETO/eR1-CreER mice, induction of RUNX1-ETO in Runx1^+^ cells at 2 weeks of age led to the development of AML with spontaneous skin inflammation (Abdallah et al., 2021). Because eR1-CreER is active in both hematopoietic cells and skin, it remained unclear whether the observed skin phenotype was a direct consequence of RUNX1-ETO expression in skin or a secondary effect resulting from hematopoietic abnormalities.

To examine the keratinocyte-intrinsic effects of RUNX1-ETO, we generated tamoxifen-inducible RUNX1-ETO/K14-CreER mice, with RUNX1-ETO expression induced only in epidermal basal keratinocytes (Figure 1a). Tamoxifen was administered at 2 weeks of age, a time window previously shown to permit the efficient induction of RUNX1-ETO-driven phenotypes (Abdallah et al., 2021). The mice were then monitored over time (Figure 1b). In RUNX1-ETO/K14-CreER mice, epidermal inflammation appeared at the anterior region of the tail and in the ears 3 weeks after induction, becoming progressively more severe by 6 weeks (Figure 1c). The affected skin exhibited a rough, dry, scaling surface. Notably, the distribution of inflammation was not uniform: although inflammation occurred in the tail, ears, and hind limbs, no overt inflammation was observed on the dorsal skin (Figure 1d). The mice also developed splenomegaly beginning approximately 3 weeks after tamoxifen administration (Figure 1e). Consistently, peripheral blood analysis revealed leukocytosis, which was significantly more pronounced at 3 weeks in comparison with control mouse white blood cell counts (Figure 1f).

**Figure 1.**
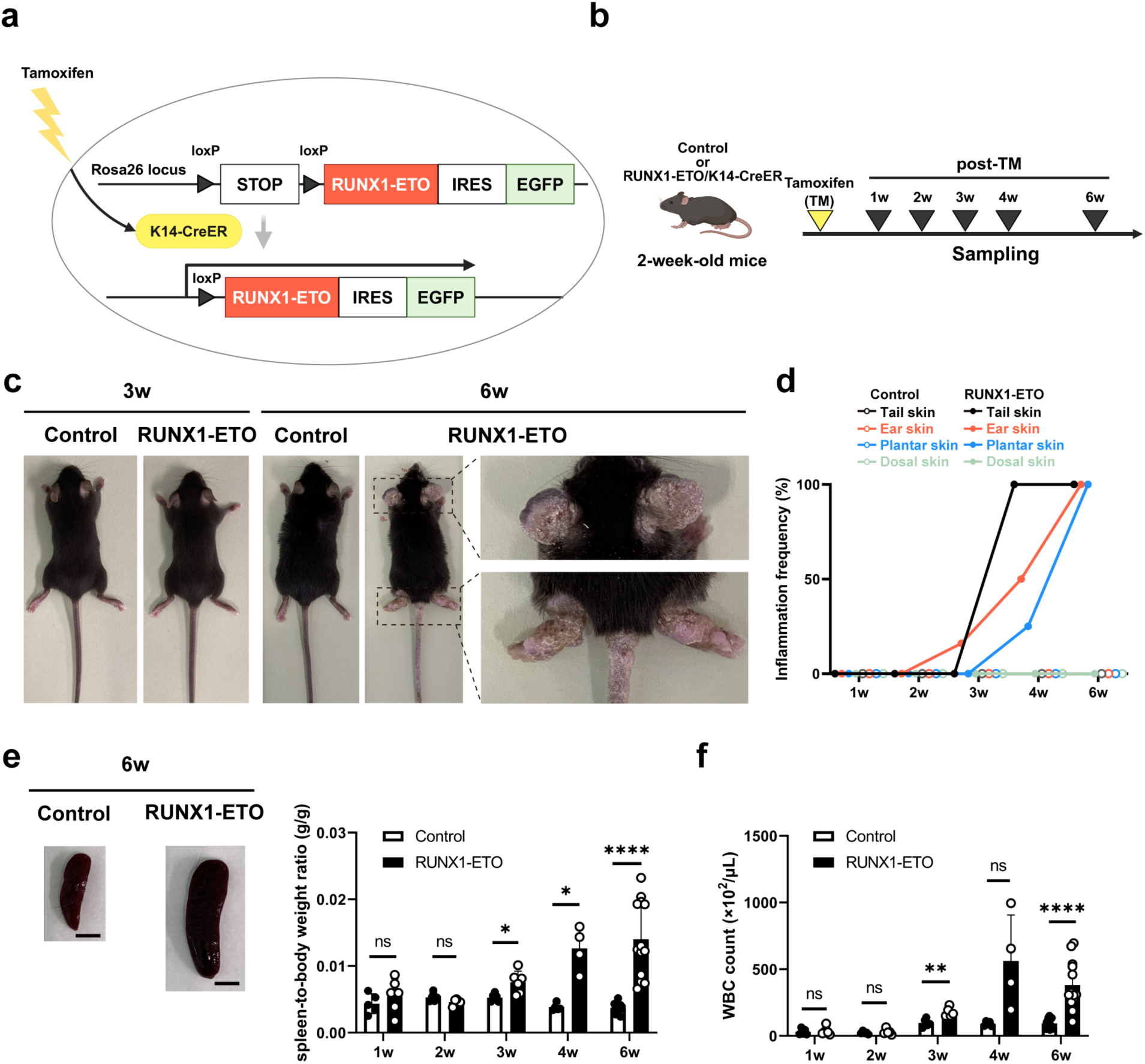
Inducible epidermal RUNX1-ETO expression is associated with progressive inflammatory pathology. **(a)** Schematic representation of the inducible RUNX1-ETO/K14-CreER construct used for inducible expression in epidermal keratinocytes. **(b)** Experimental timeline. Control or RUNX1-ETO mice received a single intraperitoneal injection of tamoxifen (TM; 100 µg/g body weight) at 2 weeks of age. Mice were sacrificed at 1, 2, 3, 4, and 6 weeks post-TM. **(c)** Representative gross appearance of control and RUNX1-ETO mice following TM administration. Insets show higher-magnification views of inflamed regions outlined by black dashed boxes. **(d)** Time course of the proportion of mice exhibiting visible skin inflammation in control or RUNX1-ETO/K14-CreER mice. **(e)** Representative images of spleens from control and RUNX1-ETO mice and quantification of spleen-to-body weight ratios. Scale bar: 5 mm. **(f)** White blood cell (WBC) counts in the peripheral blood of control and RUNX1-ETO mice. Data are presented as the mean ± SD. Each dot represents one mouse. Significance was assessed using a two-tailed unpaired Student’s *t* test. \**p* < 0.05; \*\**p* < 0.01; \*\*\*\**p* < 0.0001; ns, not significant.

Because the RUNX1-ETO cassette contains an EGFP reporter, its expression can be visualized by GFP fluorescence (Figure 1a). Whole-mount imaging of the tail epidermis was performed to assess the efficiency and distribution of RUNX1-ETO induction (Figure 2a). By 3 weeks post-induction, a clear GFP signal was observed throughout the epidermis, indicating widespread induction of RUNX1-ETO. This accumulation further increased by 6 weeks. (Figure 2b)

**Figure 2.**
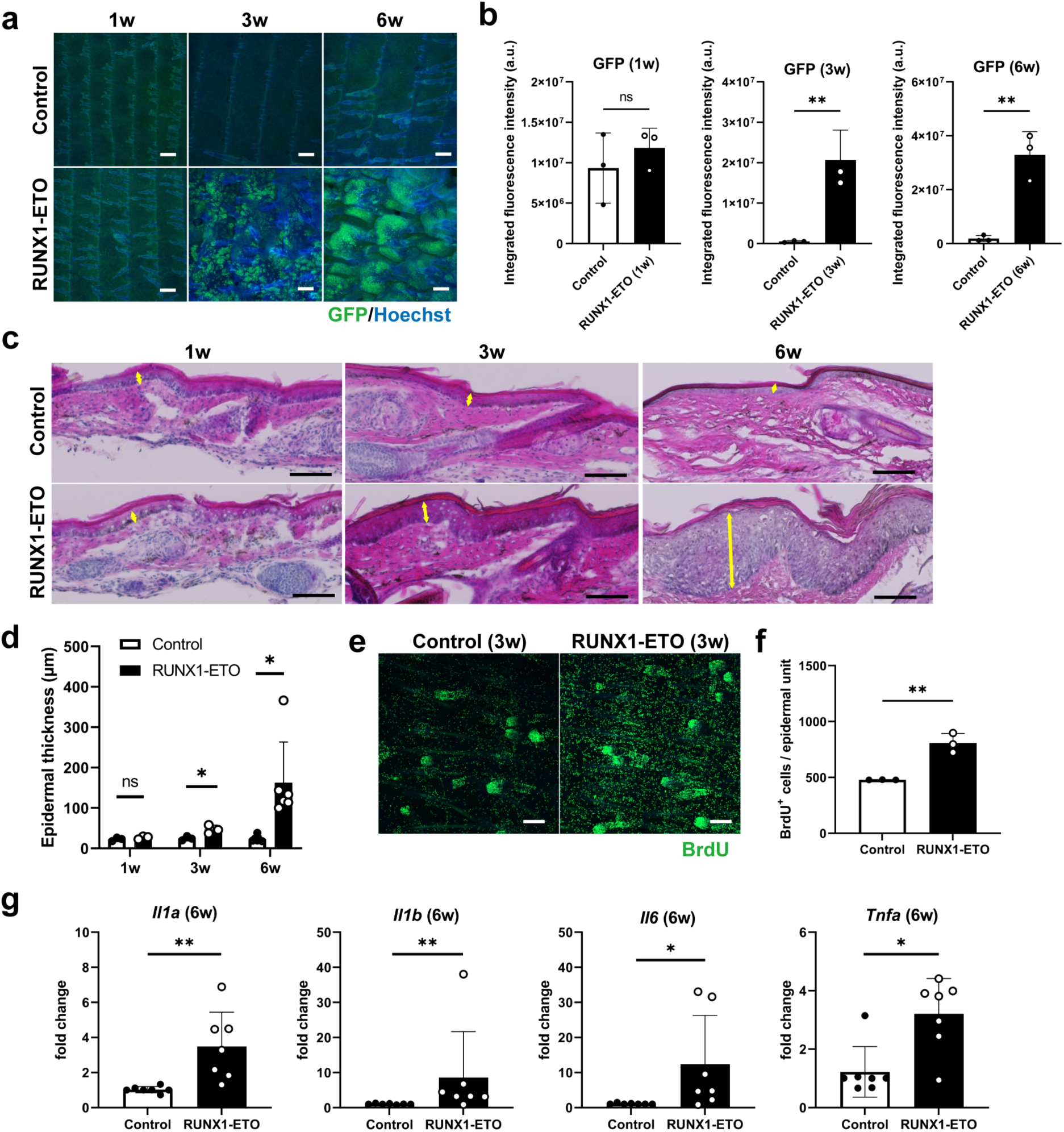
RUNX1-ETO expression induces inflammatory features in tail skin (a, **b)** Representative whole-mount GFP staining images of tail epidermis from mice at 1, 3, and 6 weeks post-TM and quantification of GFP fluorescence intensity (b). RUNX1-ETO (GFP, green); nuclei (Hoechst, blue). Scale bar: 200 µm. **(c, d)** Representative hematoxylin and eosin–stained sections of tail skin from control and RUNX1-ETO mice and quantification of epidermal thickness (d). Scale bar: 100 µm. **(e, f)** Representative whole-mount BrdU staining images of tail epidermis from mice at 3 weeks post-TM and quantification of BrdU-positive cells (f). BrdU (green); nuclei (Hoechst, blue). Scale bar: 200 µm. **(g)** Quantitative RT–PCR analysis of inflammatory cytokines in tail skin. Data are presented as the mean ± SD. Each dot represents one mouse. Significance was assessed using a two-tailed unpaired Student’s *t* test. \**p* < 0.05; \*\**p* < 0.01; ns, not significant.

A histological analysis of the tail skin revealed increased epidermal thickness as early as 3 weeks after tamoxifen induction (Figure 2c and d). To determine whether this epidermal thickening was associated with increased cell proliferation, BrdU incorporation was assessed 3 weeks after induction. Whole-mount staining revealed that epidermal proliferation was significantly increased in RUNX1-ETO mice compared with that in control mice (Figure 2e and f).

To further evaluate inflammatory changes, quantitative qRT–PCR was performed on tail skin samples collected 6 weeks after tamoxifen induction, when inflammatory changes were pronounced. The expression of multiple pro-inflammatory cytokines, including IL-1α, IL-1β, IL-6, and TNF-α, was significantly increased in RUNX1-ETO/K14-CreER mice (Figure 2g). Together, these results indicate that RUNX1-ETO expression in epidermal keratinocytes induces epidermal hyperproliferation and inflammatory activation, leading to progressive skin inflammation.

### Site-specific skin inflammation in RUNX1-ETO mice preferentially occurs in hairless regions

Inflammatory lesions in RUNX1-ETO/K14-CreER mice were preferentially observed in hairless regions, such as the tail, ears, face, and hind limbs (Figure 1c). In contrast, no overt inflammatory phenotype or hair abnormalities were observed in the hair-bearing dorsal skin. This regional difference in inflammatory phenotype suggests that mechanical stress or environmental exposure may promote inflammatory responses in susceptible skin sites in this model.

To determine whether this regional difference was also reflected at the histological level, we examined epidermal architecture in multiple skin sites. Consistent with macroscopic observations (Figure 1c, d), histological analyses of the ear (Figure 3a and b) and plantar skin (Figure 3c and d) revealed increased epidermal thickness 6 weeks after tamoxifen administration. In contrast, the dorsal skin showed no detectable increase in epidermal thickness and remained comparable to that of control mice (Figure 3e and f). The preferential occurrence of inflammation in mechanically exposed skin sites is reminiscent of the Koebner phenomenon described in inflammatory skin diseases, in which mechanical stress promotes lesion formation (Weiss et al., 2002).

**Figure 3.**
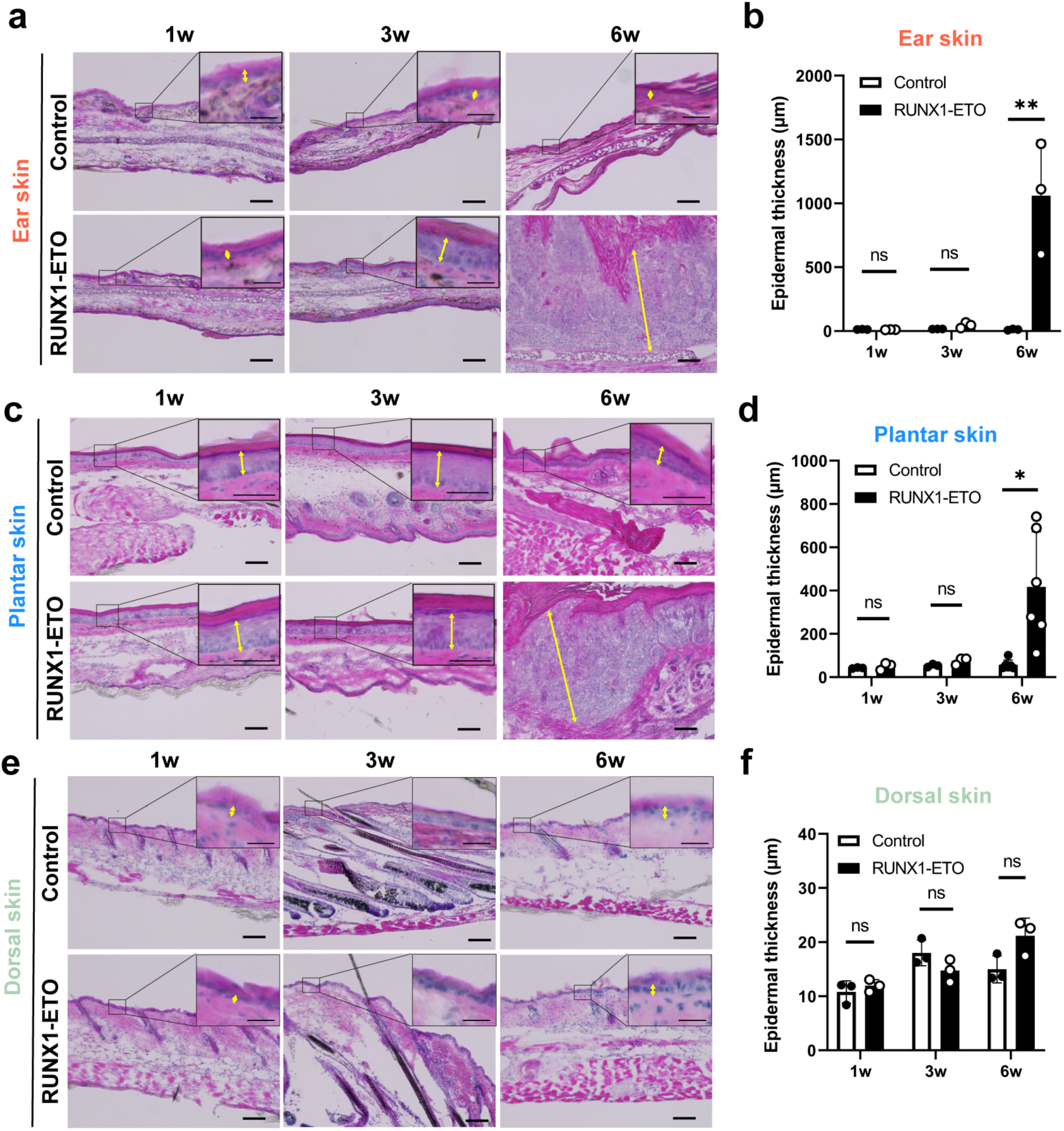
RUNX1-ETO–induced skin inflammation shows regional differences across skin sites (a–f) Representative hematoxylin and eosin–stained sections of ear skin (a, b), plantar skin (c, d), and dorsal skin (e, f) from control and RUNX1-ETO mice and quantification of epidermal thickness (b, d, f). Insets show higher-magnification views of the indicated regions. Scale bars: 100 µm; 25 μm (inset in a); 50 μm (inset in c). Data are presented as the mean ± SD. Each dot represents one mouse. Significance was assessed using a two-tailed unpaired Student’s *t* test. \**p* < 0.05; \*\**p* < 0.01; ns, not significant.

### RUNX1-ETO disrupts the spatial organization of epidermal stem cell compartments

We next analyzed how RUNX1-ETO expression affects epidermal stem cell organization. Epidermal stem cells in the tail skin are organized into slow-cycling and fast-cycling populations, which give rise to K10^+^ interscale and K36^+^ scale differentiation lineages, respectively (Sada et al., 2016) (Figure 4a). These populations form a characteristic spatial pattern in which clusters of fast-cycling lineage cells in the scale regions are regularly distributed and surrounded by interscale regions comprising slow-cycling lineage cells (Figure 4a). This spatial organization can be visualized by whole-mount staining using K36 to mark scale regions and K10 to label interscale regions.

**Figure 4.**
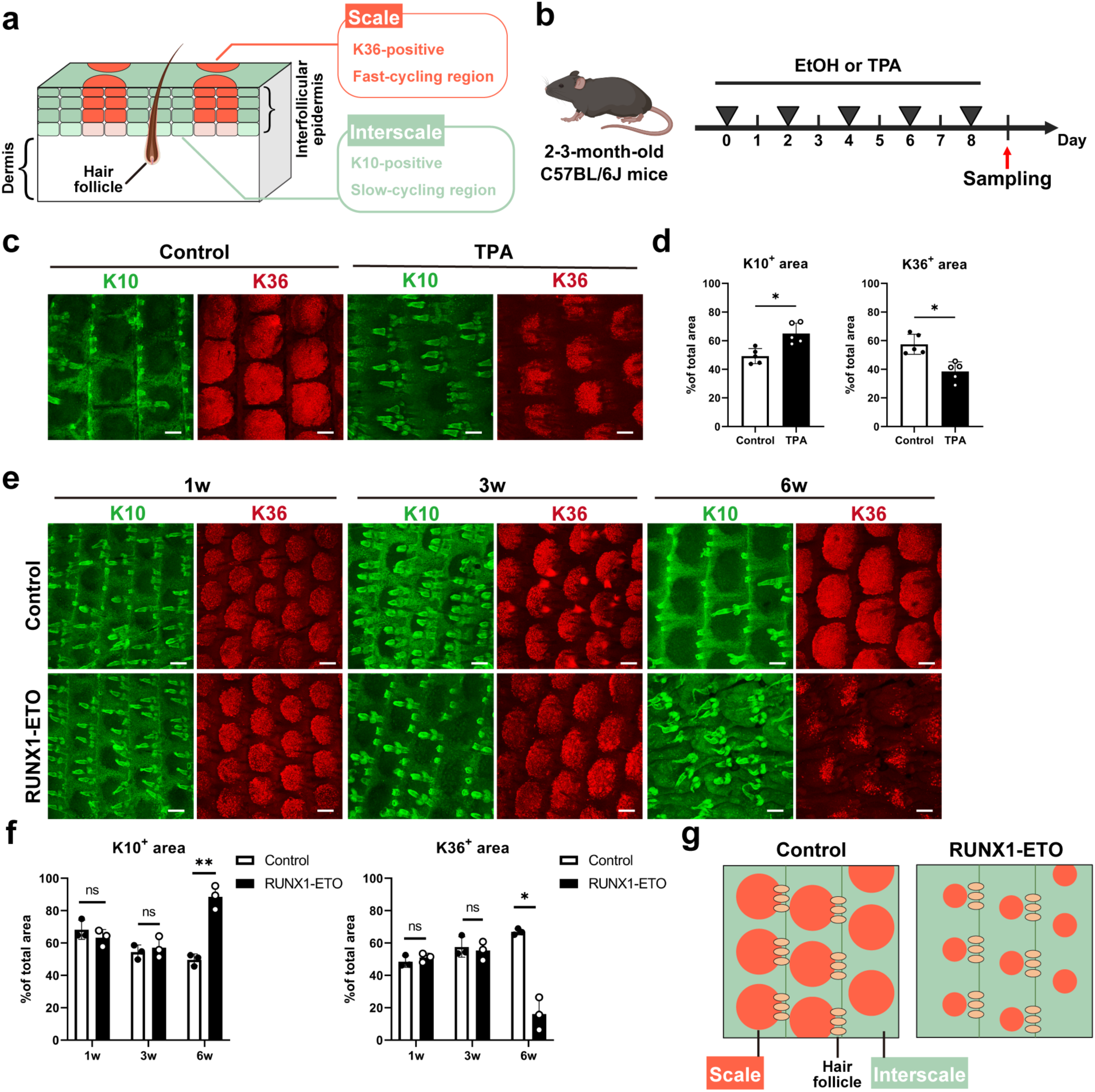
RUNX1-ETO expression impairs epidermal stem cell compartments in tail skin. **(a)** Schematic representation of scale (K36⁺, red) and interscale (K10⁺, green) regions in mouse tail skin. **(b)** Experimental timeline. C57BL/6J mice were treated with 100% ethanol (EtOH) or TPA every other day and sacrificed after five treatments. **(c, d)** Representative whole-mount immunostaining images of K10 and K36 of tail epidermis from control and TPA-treated mice and the quantification of interscale and scale areas (d). K10 (interscale, green); K36 (scale, red); nuclei (Hoechst, blue). Scale bars: 200 µm. **(e, f)** Representative whole-mount immunostaining images of K10 and K36 of tail epidermis from control and RUNX1-ETO mice and the quantification of interscale and scale areas (f). K10 (interscale, green); K36 (scale, red); nuclei (Hoechst, blue). Scale bars: 200 µm. **(g)** Schematic illustration of inflammation-associated reduction of the scale region. Data are presented as the mean ± SD. Each dot represents one mouse. Significance was assessed using a two-tailed unpaired Student’s *t* test. \**p* < 0.05; \*\**p* < 0.01; ns, not significant.

Previous studies have demonstrated that this compartmentalized structure is sensitive to chronological skin aging (Raja et al., 2022) and inflammatory stimuli (Phung et al., 2026), both of which lead to shrinkage or the disruption of scale regions. Consistent with the aforementioned findings, the topical application of 12-O-tetradecanoylphorbol-13-acetate (TPA) induced a marked reduction in the size of the scale regions in the tail epidermis (Figure 4b–d).

Notably, a similar alteration in the interscale–scale compartment was observed in RUNX1ETO/K14-CreER mice (Figure 4e). Six weeks after tamoxifen administration, the size of the K36^+^ scale regions was significantly reduced compared with that in control mice, accompanied by the expansion of K10^+^ interscale regions (Figure 4f). These results indicate that the intrinsic perturbation of keratinocytes by RUNX1-ETO disrupts the maintenance of fast-cycling stem cell compartments, leading to pathological remodeling of the spatial organization of epidermal stem cell populations (Figure 4g).

### RUNX1-ETO shifts epidermal basal keratinocytes from a homeostatic to an inflammatory gene expression state

To investigate gene expression changes underlying RUNX1-ETO-induced skin inflammation, we isolated integrin α6^high^/CD34^-^/Sca1^+^ epidermal basal keratinocytes from the tail skin of RUNX1-ETO/K14-CreER mice 3 weeks after tamoxifen induction and subjected them to RNA-seq analysis. Taking advantage of the GFP reporter co-expressed upon RUNX1-ETO induction, the cells were further gated according to GFP fluorescence to isolate RUNX1-ETO-expressing cells (Figure 5a and b). Basal keratinocytes isolated from control mice were used for comparison.

**Figure 5.**
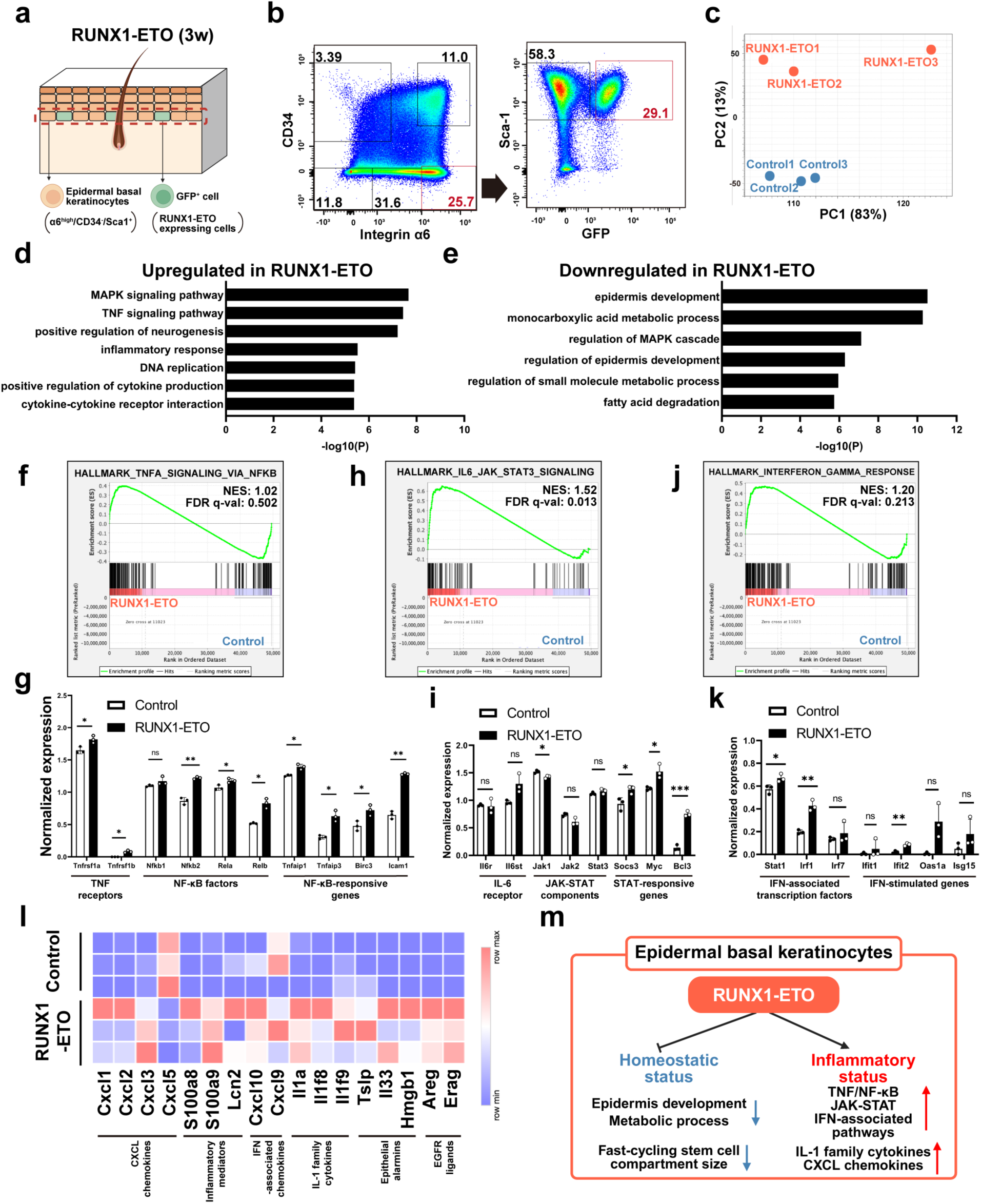
RNA-seq analysis reveals inflammatory pathway activation in RUNX1-ETO^+^ epidermal basal keratinocytes (a, **b)** Schematic representation of the tail skin and FACS gating strategy used to isolate GFP^+^ epidermal basal keratinocytes from RUNX1-ETO mice. **(c)** Principal component analysis (PCA) of RNA-seq data from epidermal basal keratinocytes of control and RUNX1-ETO mice. **(d, e)** Gene Ontology (GO) analysis of genes upregulated (d) or downregulated (e) in RUNX1-ETO^+^ epidermal basal keratinocytes compared with controls. **(f–k)** Gene set enrichment analysis (GSEA) of inflammation-related hallmark gene sets and expression plots of representative genes (g, i, k). **(l)** Heatmap showing cytokine and chemokine expression in control and RUNX1-ETO^+^ epidermal basal keratinocytes. **(m)** Summary of RUNX1-ETO–associated changes in epidermal keratinocytes, highlighting suppression of homeostatic programs and activation of inflammatory pathways. Data are presented as the mean ± SD. Each dot represents one mouse. Significance was assessed using a two-tailed unpaired Student’s *t* test. \**p* < 0.05; \*\**p* < 0.01; \*\*\**p* < 0.001; ns, not significant.

Principal component analysis (PCA) revealed a clear separation between RUNX1-ETO and controls, indicating global changes in gene expression profiles in basal keratinocytes (Figure 5c). Gene Ontology (GO) analysis of upregulated genes revealed enrichment in pathways related to inflammatory signaling, including the MAPK and TNF pathways, as well as cytokine production (Figure 5d and Supplemental Table S1). Genes related to DNA replication and cell cycle regulation were also enriched, suggesting that RUNX1-ETO expression is accompanied by increased proliferative activity in basal keratinocytes.

In contrast, downregulated genes were enriched in pathways related to epidermal development and fatty acid metabolism (Figure 5e and Supplemental Table S1). Consistent with this, Runx1 has been shown to influence the lipid metabolism-related genes, such as *Scd1* and *Soat1*, in hair follicle stem cells (Jain et al., 2018). These results suggest that RUNX1-ETO induction impairs homeostatic gene expression programs in basal epidermal keratinocytes and shifts these cells toward an inflammatory and proliferative state. These transcriptional changes are consistent with the epidermal hyperproliferation and inflammatory phenotypes observed in RUNX1-ETO/K14-CreER mice.

### RUNX1-ETO activates inflammatory gene programs in epidermal basal keratinocytes

We performed gene set enrichment analysis (GSEA) to further characterize inflammatory pathways activated by RUNX1-ETO. Among inflammatory pathways, gene sets related to TNFα/NF-κB and IL-6–JAK–STAT3 signaling were enriched in RUNX1-ETO^+^ basal keratinocytes (Figure 5f, h). Consistent with these enrichments, RUNX1-ETO^+^ basal keratinocytes showed increased expression of TNF receptors (*Tnfrsf1a* (TNFR1) and *Tnfrsf1b* (TNFR2)), NF-κB-associated transcription factors (*Nfkb2*, *Rela*, and *Relb*), TNF/NF-κB–responsive genes (*Tnfaip1*, *Tnfaip3*, *Birc3*, and *Icam1*) (Figure 5g), and STAT-associated genes (*Socs3*, *Myc*, and *Bcl3*) (Figure 5i). In addition, the expression of several interferon-responsive genes, including *Stat1*, *Irf1*, and *Ifit2*, was significantly upregulated (Figure 5j and k).

An analysis of inflammatory mediators further revealed increased expression of keratinocyte-derived IL-1 family cytokines, including *Il1a*, *Il1f8* (*Il36b*), and *Il1f9* (*Il36g*) (Figure 5l). In addition, the expression of multiple CXCR2 ligands, including *Cxcl1*, *Cxcl2*, *Cxcl3*, and *Cxcl5*, was also upregulated, together with inflammatory mediators (*S100a8*, *S100a9*, and *Lcn2*) and interferon-associated chemokines (*Cxcl10* and *Cxcl9*). Together, these results indicate that RUNX1-ETO expression in epidermal basal keratinocytes is associated with inflammatory programs involving NF-κB–associated signaling, STAT-related responses, and interferon-stimulated gene expression (Figure 5m). These changes are consistent with the increased expression of inflammatory cytokines observed in the whole skin at later stages (Figure 2g).

## Discussion

The current study demonstrates that the disruption of Runx1-dependent regulatory programs in epidermal basal keratinocytes can impair epidermal homeostasis and induce a progressive inflammatory skin phenotype *in vivo*. Epidermal keratinocytes function not only as a physical barrier but also as active regulators of skin immune responses; however, how perturbations of keratinocyte cellular state initiate inflammatory skin pathology remains poorly understood. Our findings support the concept that keratinocyte-intrinsic transcriptional dysregulation can drive inflammatory processes, rather than merely respond to immune cell-derived signals. Although RUNX1-ETO expression was restricted to epidermal keratinocytes in our model, the mice developed systemic inflammatory changes, including splenomegaly and leukocytosis. Such systemic responses are commonly observed in inflammatory skin conditions and may reflect secondary cytokine signaling originating from inflamed epidermal tissue. Notably, our RUNX1-ETO model preferentially developed skin inflammation in hairless skin regions, whereas hair-bearing dorsal skin remained largely unaffected. This regional susceptibility may reflect differences in mechanical or environmental exposure, reminiscent of the Koebner phenomenon described in inflammatory skin diseases (Weiss et al., 2002). Hairless skin is more directly exposed to mechanical stress, which may exacerbate inflammatory signaling in RUNX1-ETO–expressing keratinocytes. Thus, this model provides a useful genetic system for studying how keratinocyte-intrinsic transcriptional perturbations interact with mechanical or environmental factors, thereby promoting inflammatory skin pathology. This mechanism may be relevant to inflammatory skin diseases in which mechanical stress and keratinocyte-intrinsic signaling contribute to lesion development, such as psoriasis or other Koebner-associated conditions.

Runx1 has been implicated in the regulation of hair follicle stem cell proliferation, in part by suppressing transcription of cell cycle inhibitors (Hoi et al., 2010; Lee et al., 2013; Osorio et al., 2008). Consistent with this role, RUNX1-ETO-expressing keratinocytes exhibited increased proliferative activity and cell-cycle-related gene expression. However, the strong induction of inflammatory gene expression programs indicates that the phenotype cannot be explained solely by hyperproliferation but rather reflects inflammatory remodeling of the epidermis.

The alteration of epidermal stem cell compartments observed in the RUNX1-ETO model (Figure 4e, f) suggests that keratinocyte-intrinsic perturbations can compromise epidermal spatial organization. Inflammatory signaling has been shown to remodel epidermal stem cell dynamics, in part through the IL-1–mediated suppression of Wnt activity (Phung et al., 2026), suggesting that similar mechanisms may contribute to the phenotype observed here.

The increased expression of IL-1 family cytokines and CXCL chemokines observed in the RUNX1-ETO model is consistent with the involvement of multiple inflammatory transcriptional programs in keratinocytes, including the TNF/NF-κB, STAT, and interferon-associated pathways. Notably, *Il1a* expression was upregulated in RUNX1-ETO basal keratinocytes at 3 weeks post-induction, whereas *Il1b* expression was increased in the whole tail skin at a later stage, when skin inflammatory phenotypes became more prominent. This finding raises the possibility that IL-1α is produced primarily by RUNX1-ETO–expressing keratinocytes, whereas IL-1β may originate from other cell populations in the inflamed skin, possibly including myeloid cells. This observation is consistent with the findings of previous reports that have demonstrated that keratinocyte-derived alarm signals, including IL-1α, can promote epidermal hyperproliferation and inflammatory cell recruitment, whereas myeloid cell-derived IL-1β may further amplify IL-1 signaling during skin inflammation (Macleod et al., 2021; Morizane et al., 2023). Thus, RUNX1-ETO expression in basal keratinocytes may initiate an inflammatory response that is subsequently reinforced by crosstalk with immune cells.

Previous studies have revealed a functional link between Runx1 and NF-κB signaling, although the direction of regulation appears to depend on tissue context. For example, Runx1 suppresses NF-κB activation by interacting with IKKβ in lung epithelial cells (Tang et al., 2017) and modulating the IKK complex in hematopoietic cells (Nakagawa et al., 2011). In contrast, Runx1 enhances NF-κB activity by interacting with the p50 subunit in macrophages (Luo et al., 2016), and cooperates with NF-κB and AP-1 transcription factors to induce the expression of inflammatory cytokines, such as IL-23, in intestinal epithelial cells (Lim et al., 2020). In skin cancer, Runx1 has been implicated in Stat3 signaling as it regulates Socs family members (Scheitz et al., 2012). Moreover, RUNX1-ETO has previously been shown to induce a type I interferon response in leukemia (DeKelver et al., 2014), suggesting that activation of interferon-associated transcriptional changes may represent a conserved response to RUNX1-ETO. Together, these findings suggest that the disruption of Runx1-dependent transcriptional regulation by RUNX1-ETO may shift basal keratinocytes toward a pro-inflammatory transcriptional state through the coordinated activation of the NF-κB-, STAT-, and interferon-associated pathways.

An additional important consideration is that the prominent inflammatory phenotype described here has not generally been reported in the skin of previously characterized Runx1 mutant models, including RUNX1-flox/K14-CreER mice (Hoi et al., 2010). This discrepancy suggests that the phenotype cannot be explained solely by Runx1 loss of function. Instead, transcriptional dysregulation that is unique to the RUNX1-ETO fusion protein, such as dominant-negative activity combined with chromatin remodeling or altered transcription factor recruitment (Ptasinska et al., 2012), may contribute to the phenotype.

Several limitations should be acknowledged. First, we have not identified the direct genomic binding sites or transcriptional targets of RUNX1-ETO in epidermal keratinocytes. Future studies combining ChIP-seq or CUT&RUN with chromatin accessibility analyses will be required to determine whether induction of the IL-1/IL-36-CXCL axis is directly regulated by RUNX1-ETO. Second, this model does not recapitulate a known disease-causing mutation in human skin; rather, it represents an experimentally induced perturbation. Accordingly, the current data should be interpreted as a broader principle that disruption of transcriptional control in basal keratinocytes can initiate or amplify inflammatory conditions, rather than directly modeling a specific skin disease.

Together, our findings identify Runx1-dependent transcriptional regulation as an important mechanism for maintaining epidermal immune homeostasis and suggest that its disruption in keratinocytes can initiate inflammatory skin pathology.

## Materials and Methods

### Mice

The mice were maintained under specific pathogen-free conditions at the Center for Animal Resources and Development (CARD) at Kumamoto University and at the Laboratory of Embryonic and Genetic Engineering at Kyushu University. The mice were maintained on a 12-h light/12-h dark cycle at a controlled temperature, with *ad libitum* access to water and standard chow. All experimental procedures were conducted in accordance with the guidelines for animal experimentation approved by the Institutional Animal Experiment Committee of Kumamoto University and Kyushu University. C57BL/6J mice were purchased from Charles River Laboratories Japan, Inc. (Kanagawa, Japan) and Japan SLC, Inc. (Shizuoka, Japan). The generation of RUNX1-ETO mice (CARD ID: 2896; C57BL/6J background) has been described previously (Abdallah et al., 2021). RUNX1-ETO mice were crossed with K14-CreER mice (The Jackson Laboratory, strain 005107; C57BL/6J background) to obtain double transgenic mice. Both male and female mice were used for the experiments.

### Tamoxifen administration

Tamoxifen (Sigma-Aldrich) was dissolved in corn oil (Sigma-Aldrich) by heating at 70°C. A single intraperitoneal injection of tamoxifen (100 μg/g body weight) was administered to 2-week-old RUNX1-ETO/K14-CreER mice. Mice were collected and analyzed at 1, 2, 3, 4, and 6 weeks post-tamoxifen administration.

### BrdU administration

BrdU (5-bromo-2′-deoxyuridine; Sigma-Aldrich) was administered by intraperitoneal injection (2.5 μg/μL) once daily for two consecutive days before sacrifice.

### TPA-induced skin inflammation

To induce local skin inflammation, 12-O-tetradecanoylphorbol-13-acetate (TPA; Sigma-Aldrich) was applied to the whole tail skin of the mice (10 μg in 40 μL of 100% ethanol) at 2-day intervals for a total of five applications. Control mice received an equivalent volume of vehicle (100% ethanol). Skin samples were collected 1 day after the final application.

### Sample collection and analysis

Tail, ear, plantar, and dorsal skin was collected, along with the spleen and blood, from control or RUNX1-ETO/K14-CreER mice at the indicated time points after tamoxifen induction. Spleens were weighed immediately after collection. Blood samples were collected from the inferior vena cava 1 and 2 weeks after tamoxifen induction, and from the retro-orbital sinus at 3, 4, and 6 weeks after tamoxifen induction. White blood cell counts were measured using a hematology analyzer (Celltac-a MEK-6550, Nihon Kohden). Tail skin was collected from TPA-treated mice for analysis.

### Hematoxylin and eosin staining of skin sections

Tail, ear, and dorsal skin samples were embedded in optimal cutting temperature (OCT) compound (Tissue-Tek, Sakura). Cryosections (10 µm) were cut using a cryostat and fixed in 4% paraformaldehyde for 10 min at room temperature. Sections were stained with hematoxylin (Wako) for 1 min 30 s and eosin Y (Wako) for 15 s. Images were acquired using an EVOS M5000 imaging system (Thermo Fisher Scientific).

### Whole-mount immunostaining and imaging

Whole-mount immunostaining of tail skin was performed as previously described (Sada et al., 2016). The following primary antibodies were used: rat anti-BrdU (1:300; Abcam, ab6326), mouse anti-K10 (1:100; Abcam, ab9026), guinea pig anti-K36 (1:100; Proteintech, 14309-1-AP), and rat anti-GFP (1:100; Abcam, ab13970). For BrdU detection, samples were treated with 2 N hydrochloric acid at 37 °C for 30 min before antibody incubation. Nuclei were stained with Hoechst dye (Sigma–Aldrich). Epidermal specimens were imaged using confocal microscopy (Nikon A1 HD25 or Zeiss LSM900). Images were acquired and processed using NIS-Elements Imaging or ZEN (Blue Edition) software, with brightness and contrast adjustments performed in Adobe Photoshop (version 2024). All imaging parameters and post-acquisition processing settings were kept constant between control and experimental samples. All whole-mount images are presented as Z-stack maximum projections viewed from the basal layer.

### FACS isolation

Epidermal cell isolation and sorting were performed as previously described (Sada et al., 2016) with minor modifications. Briefly, subcutaneous tissues were removed from the excised tail skin, followed by incubation in 0.25% trypsin/EDTA at 4 °C overnight. After an additional 30 min of incubation at 37 °C, the epidermal keratinocytes were gently dissociated to obtain a single-cell suspension.

The cells were stained on ice for 30 min with the following antibodies: biotin-conjugated anti-CD34 (1:50; eBioscience, 13-0341), APC-conjugated streptavidin (1:100; BD Biosciences, 554067), BV510-conjugated integrin α6 (1:200; BD Biosciences, 563271), and BV421-conjugated Sca-1 (1:100; BD Biosciences, 562729). Dead cells were excluded using 7-AAD (BD Biosciences).

Epidermal basal keratinocytes were defined as integrin α6^high^/CD34^−^/Sca-1^+^ cells. In RUNX1-ETO mice, GFP^+^ cells within this population were sorted, whereas in wild-type mice, the entire basal cell population was collected. Cell sorting was performed using a FACSAria flow cytometer (BD Biosciences), and data were analyzed with FlowJo software (BD Biosciences).

### RNA-sequencing

FACS-isolated epidermal basal keratinocytes from wild-type and RUNX1-ETO/K14-CreER mice were stored in TRIzol reagent (Ambion, 10296028). Approximately 1 × 10⁶ cells were collected per sample, and three biological replicates were analyzed for each group. RNA extraction, library preparation, and sequencing were performed at Tsukuba i-Laboratory LLP. RNA integrity was assessed using an Agilent 2100 Bioanalyzer. RNA-seq libraries were prepared using the SMARTer Stranded Total RNA-Seq Kit v2–Pico Input Mammalian (Takara) and sequenced on an Illumina NextSeq 500 platform.

Raw sequencing data were processed and analyzed using CLC Genomics Workbench (Qiagen). Gene expression levels were normalized using the reads per kilobase per million mapped reads (RPKM) method. Differential expression analysis was performed with Benjamini–Hochberg correction to control the false discovery rate, and genes with an absolute log_2_ fold change greater than 2 (|log_2_FC| > 2) were selected.

PCA was performed in CLC Genomics Workbench using genes with the highest variance across samples. GO analysis was conducted using Metascape. GSEA was performed using GSEA software with the MSigDB Hallmark gene sets. The genes shown in the graphs and heatmaps were selected according to differential expression and biological relevance. Normalized expression values of selected genes were used for statistical analyses and visualization in GraphPad Prism 9. Heatmaps were generated using MORPHEUS according to normalized expression values obtained from CLC Genomics Workbench (Qiagen).

### Quantitative RT–PCR

Total RNA was extracted from tail skin using an RNeasy Mini Kit (Qiagen) according to the manufacturer’s directions. cDNA was synthesized using iScript Reverse Transcription Supermix (Bio-Rad). Quantitative RT–PCR was performed on a LightCycler 96 system (Roche) with FastStart Essential DNA Green Master (Roche), using 5 ng of cDNA per reaction. Relative gene expression levels were calculated using the 2^−ΔΔCt^ method.

The primer sequences used were as follows:

*Il1a* forward (F) 5’-CTGGAAGAGACCATCCAACCC-3’ and reverse (R) 5’-ACTTCCTGTTGCAGGTCATTT-3’; *Il1b* F 5’-AAGAGCCCATCCTCTGTGACT-3’ and R 5’-GGAGAATATCACTTGTTGGTT-3’; *Il6* F 5’-CCAGTTGCCTTCTTGGGACTG-3’ and R 5’-GCCATTGCACAACTCTTTTCTC-3’; *Tnfa* F 5’-GCCCACGTCGTAGCAAACCAC-3’ and R 5’-GCAGGGGCTCTTGACGGCAG-3’; *Actb* F 5’-GATCAGCAAGCAGGAGTACGA-3’ and R 5’-AAAACGCAGCTCAGTAACAGTC-3’.

### Quantification and statistical analysis

All quantifications were conducted with at least three independent experiments. Data are presented as the mean ± standard deviation (SD). GraphPad Prism 9 (GraphPad Software) was used for statistical analyses; normality was assessed using the Shapiro–Wilk test, when applicable. For comparisons between two groups, two-tailed unpaired Student’s *t* tests were used. A *p* value < 0.05 was considered to indicate significance. Sample sizes were determined according to experimental considerations. Experiments were not randomized, and investigators were not blinded to group allocation or outcome assessment.

For histological quantification, epidermal thickness was measured according to previously defined epidermal units (Changarathil et al., 2019), with at least six units analyzed per mouse. BrdU-positive cells were manually counted from 3–4 projected Z-stack images per mouse, each containing 7–15 scale or interscale structures. K10- and K36-positive areas were quantified from 6–8 projected Z-stack images per mouse, each containing approximately 15 scale or interscale structures.

## Ethics statement

All animal experiments were reviewed and approved by the Animal Care and Use Committee of Kumamoto University and the Animal Experiment Committee of Kyushu University. The experiments were conducted in accordance with relevant institutional guidelines for animal research.

## Data availability statement

The RNA sequencing datasets obtained in this study have been deposited in the National Center for Biotechnology Information-Gene Expression Omnibus (GEO) under accession number GSE309298 (https://www.ncbi.nlm.nih.gov/geo/query/acc.cgi?acc=GSE309298).

## Declaration of generative AI and AI-assisted technologies in the writing process

During the preparation of this manuscript, the authors used ChatGPT to assist with language clarity and readability. All content was critically reviewed and revised by the authors, who take full responsibility for the accuracy and integrity of the final manuscript.

## Acknowledgments

We thank the Center for Animal Resources and Development (CARD) at Kumamoto University, the Laboratory of Embryonic and Genetic Engineering at Kyushu University, and T. Keida (Kumamoto University) and N. Imai (Kyushu University) for mouse care. We thank the International Core Facility of Advanced Life Science at Kumamoto University and the Research Promotion Unit at Kyushu University for their support of core facilities. We thank Dr. S. Fukushima, Dr. K. Iwamoto, Dr. H. Takizawa, Dr. M. Bundo, Dr. Y. Nakachi, and Dr. S. Fujii (Kumamoto University) for helpful discussions and their critical feedback on this work. This work was supported by the Japan Agency for Medical Research and Development (AMED) under grant number JP26bm1123052 (Project for Regenerative, Cellular Medicine and Gene Therapies; to A.S.); Grant-in-Aid for Scientific Research (B) from the Japan Society for the Promotion of Science (JSPS; JP20H03266 and JP24K02035, to A.S.); Uehara Memorial Foundation (to A.S.); The Takeda Science Foundation (to A.S.); The Sumitomo Foundation (to A.S.), The Mochida Memorial Foundation for Medical and Pharmaceutical Research (to A.S.), and the Lydia O’Leary Memorial Pias Dermatological Foundation (to A.S.). This work was partly supported by the MEXT Cooperative Research Project Program, Medical Research Center Initiative for High Depth Omics, and CURE (JPMXP1323015486) to the Medical Institute of Bioregulation (MIB), Kyushu University. The infrastructure of Omics Science Center Secure Information Analysis System (OASIS; https://sis.bioreg.kyushu-u.ac.jp/), Medical Institute of Bioregulation at Kyushu University, provided some of the computational resources. We acknowledge scholarship support from a SPRING from the Japan Science and Technology Agency (JST), JPMJSP2136 (to I.N.) and the Ministry of Education, Culture, Sports, Science, and Technology-Japan (MEXT) (to H.M.P.).

## CrediT Statement (Author Contributions)

Conceptualization: A.S. and M.O.; Investigation, Formal Analysis, Validation: H.D., M.I., I.N., H.M.P., and N.T.K.N.; Methodology and Resources: M.O., A.S., and G.S.; Visualization: H.D. and A.S.; Writing—original draft preparation: H.D.; Writing—review and editing: A.S., M.I., G.S., and M.O.; Supervision: A.S.; Funding Acquisition: A.S.

